# Action Sequences Structure Human Memory Reactivation

**DOI:** 10.64898/2026.08.08.742825

**Authors:** Julia K. Schaefer, Aditya Chowdhury, Tobias Staudigl

## Abstract

Episodic memory formation is not a passive process but emerges from sequences of actions through which we engage with our environment. Yet, how these action sequences interact with the neural dynamics underlying human memory formation and retrieval remains poorly understood. Here, we investigated whether coordinated gaze shifts, one of the major exploratory actions in humans, structure the reactivation of memories. Participants learned multiple action sequences by pairing images with distinct real-world gaze shifts while we simultaneously recorded EEG and motion tracking data. During subsequent retrieval, the recalled gaze sequences acted as a direct behavioral readout of memory performance. Representational similarity analyses revealed that the reactivation of image memories was precisely timed to recalled gaze shifts within each sequence. Crucially, only correctly executed, but not incorrectly executed gaze shifts elicited significant reactivation of the corresponding image class, directly linking neural reactivation to retrieval success. These findings suggest that exploratory actions are fundamental components of episodic memories, not incidental accompaniments: self-initiated movements can serve as temporal anchors of memory reactivation that predict retrieval success. By investigating memory through natural behavior, our work provides an ecologically grounded perspective on human cognition, revealing that species-typical actions such as gaze shifts are essential to, rather than obstacles for, understanding memory’s neural basis.

## Introduction

Natural exploratory behavior in humans primarily involves coordinated movement of the head and eyes, commonly referred to as gaze (Einhäuser et al., 2007). In everyday life, gaze shifts occur across a broad range of spatial and temporal scales, from small saccades spanning a few degrees of visual angle, to large-scale gaze shifts involving coordinated head-and-eye movement (Franchak et al., 2021; Freedman, 2008). These natural gaze behaviors are not only central to how humans perceive and interact with their visual environment (Leszczynski et al., 2025; Leszczynski & Schroeder, 2019; Schroeder et al., 2010), but also influence how we encode them into memory (Flick et al., 2026; Kragel et al., 2020; Staudigl et al., 2017; Wu et al., 2025). Our memory of events is thus not simply “what we saw” but rather sequences of “what we did” and “what we saw as a consequence”. In episodic memory formation, the brain uses this sequential organization to store information about both action (“what we did”) and content (“what we saw as a consequence”) and importantly, also preserves the spatiotemporal nature of these events (Buzsáki, 2019; Buzsáki & Moser, 2013; Buzsáki & Tingley, 2018).

Despite the crucial importance of exploratory behavior, the majority of studies investigating human memory conceptualize the memory system as a passive observer, with participants encoding stimuli by viewing static images or video clips on a screen, typically while attempting to maintain fixation. Over the last years, studies have started to go beyond such fixational paradigms and are beginning to unravel how actions organize brain activity in humans, with special focus on saccades (Hoffman et al., 2013; Katz et al., 2020, 2022; Leszczynski & Schroeder, 2019; Staudigl et al., 2022) and head turns (Aghajan et al., 2017; Griffiths et al., 2024; Seeber et al., 2025). Indeed, recent studies in humans and non-human primates have begun to reveal how such interactions between neural activity and actions predict memory performance (Flick et al., 2026; Jutras et al., 2013; Kragel et al., 2020, 2021; Z.-X. Liu et al., 2020; Nikolaev et al., 2023; Popov & Staudigl, 2023; Staudigl et al., 2017; Wu et al., 2025). Further evidence that our exploratory behavior is encoded together with the sensory experience, comes from studies which show that eye-movement patterns during encoding are reinstated during successful recall (Johansson et al., 2022; Johansson & Johansson, 2014; Kragel & Voss, 2021, 2022; Nau et al., 2025; Wynn et al., 2019). However, despite these advances, a conceptual distinction is often made between “action” and “episodic memory”. An alternative account proposes that action sequences are the fundamental building blocks of memory, implying that the neural processes underlying memory formation and those implementing actions, and their sensory consequences are highly intertwined (Buzsáki et al., 2014; Nau et al., 2025; Zutshi et al., 2025).

In this work, we jointly investigated the temporal dynamics of memory for action sequences and their sensory consequences. Specifically, by integrating gaze into memory sequences, we use humans’ primary natural exploratory behavior to provide insights on how and when memory processes unfold on the neural level. We developed a novel memory paradigm in which participants were trained to encode action sequences consisting of visual stimuli and self-initiated gaze shifts. We hypothesized that the neural reactivation of their memory is tightly aligned with the reinstatement of their exploratory actions. Our primary goal was thus to measure the temporal unfolding of memory reactivation in the brain, relative to the temporal unfolding of self-paced gaze shifts. By embedding real-world head movements into the memory trace, our study offers an integrated perspective on the interplay between action sequences and human memory.

## Results

Our experimental setup consisted of five different screens where visual stimuli were presented, one directly in front of the participant, and two on each side, placed at viewing angles of 30° and 60° (see Fig.1, and Methods for details). We measured simultaneous scalp EEG, head position, and eye position while participants, without any head-restraint, performed various parts of the experiment (see Methods). In the action sequence learning part of the experiment (Fig.1a & Supplementary Material Video S.1a), participants learned 24 unique sequences of head turns. Each sequence consisted of four head turns, one to each of the four peripheral screens. Further, each such sequence was associated with four unique images, one displayed after the other, on the screens of a particular sequence. The images were drawn from four classes (faces, objects, houses, animals) presented in random order. After a distractor task, participants were asked to recall these sequences by reproducing the head-turn sequences (action sequence retrieval, Fig.1b & Supplementary Material Video S.1b). Note that the action sequence retrieval task was self-paced and that all the five screens displayed central fixation crosses only. Subsequently, participants were asked to recognize the correct images of a given sequence and sort them in the correct order onto placeholders on the central screen (image sequence sorting, Fig.1d).

**Fig. 1.**
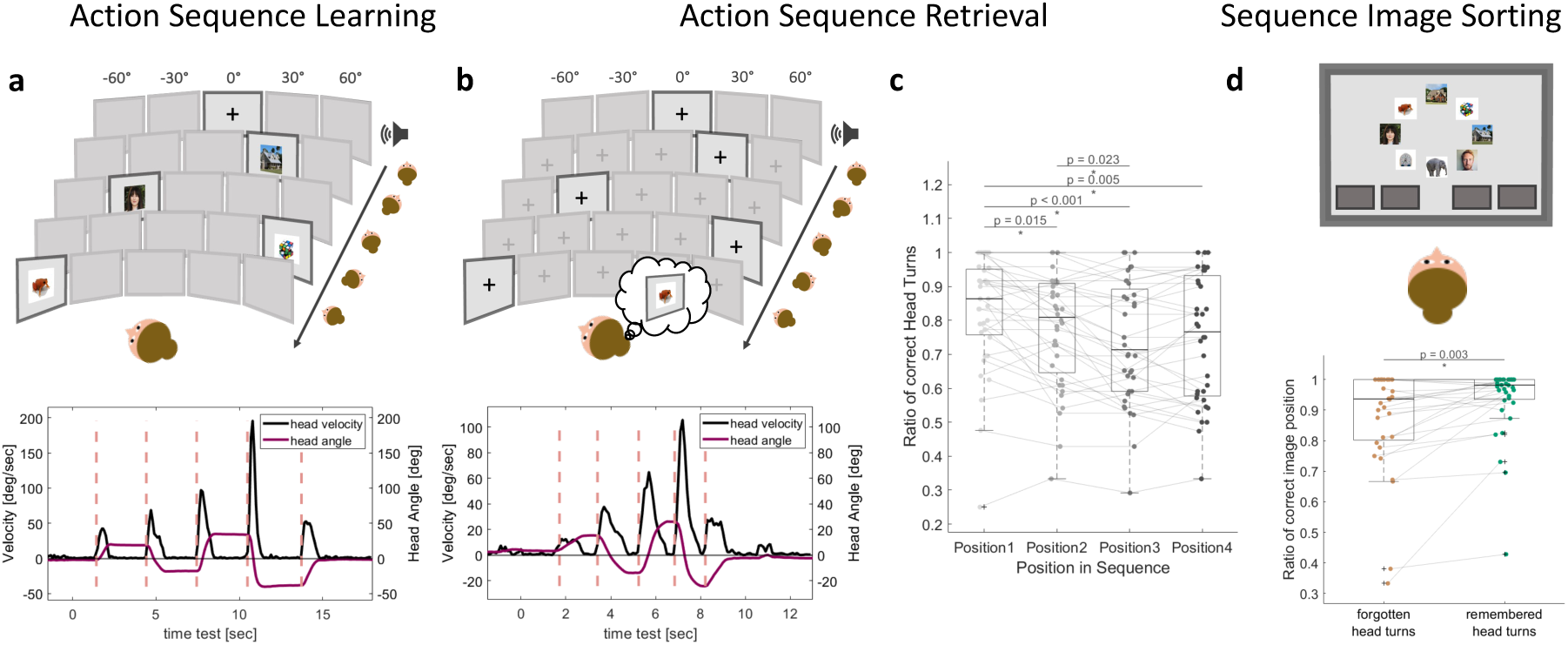
Schematic illustration of task procedure, head motion traces and behavioral performance. **a,** Illustration of the action sequence learning task, during which an auditory word cue was followed by the sequential presentation of four images, each representing one of four stimulus classes (faces, objects, houses and animals). Images appeared on four screens arranged in a horizontal semicircle at angular positions between −60° and +60° relative to the participant. Participants oriented their head toward each screen as a new image appeared, thereby learning the spatial–temporal sequence. Panel **b** illustrates the action sequence retrieval task, in which participants reproduced the four previously learned head turns while recalling the image sequence in a self-paced manner. Note that they were not asked for reports, verbal or otherwise, of their image recall during this task. The lower panels show exemplary head motion traces for **a,** action sequence learning and **b,** action sequence retrieval, for the same sequence (one participant). The dark purple line represents head angle (positive values indicate rightward head motion; negative values indicate leftward head motion). The black line indicates absolute head velocity. Head movement onsets are indicated by the rose dashed line. Each sequence was segmented according to head turn onsets into four segments. **c,** Behavioral performance across participants in the action sequence retrieval task, measured as the proportion of head turns to the correct screen, for each sequential position. All significant pair-wise comparisons, after bonferroni correction, are indicated in the panel. **d,** In the image-sorting task, participants selected the four correct images from a set of eight and placed them onto the corresponding screens in the correct sequential order. For each participant, we identified remembered and forgotten segments in the action sequence retrieval task (based on the head turn to the correct or incorrect screen) and computed, for each group, the proportion of correctly ordered images on screens in the image sorting task. We find that correctly reenacted head-turns are associated with a significantly higher performance in the image sorting task (p = 0.003). Photos: unsplash.com.

**Fig. 2.**
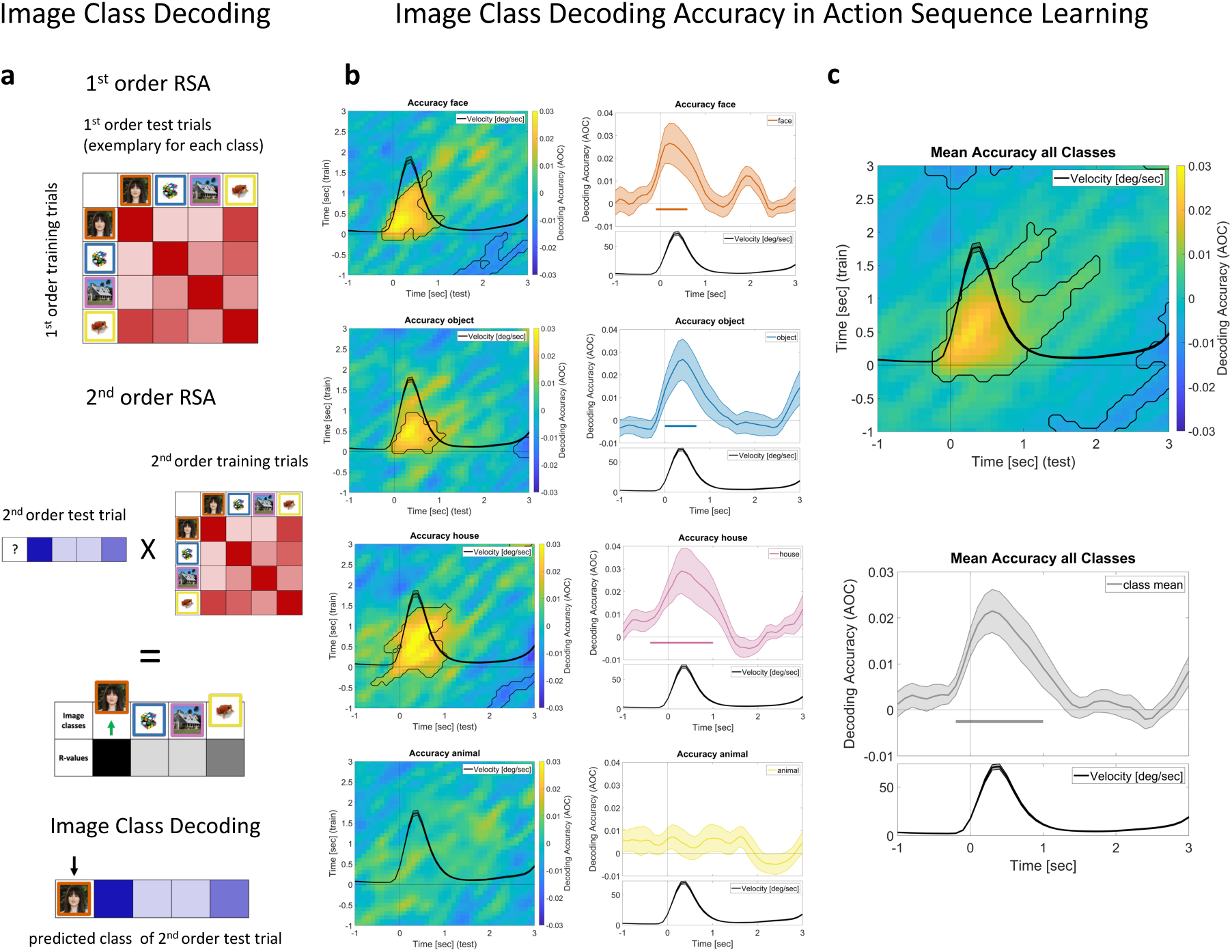
Schematic illustration of RSA based decoding approach and image class decoding accuracy during action sequence learning. **a,** Schematic illustration of analysis method for a single test and train timepoint. *First-order RSA*: Stimuli consisted of four image classes: faces (orange), objects (blue), houses (purple) and animals (yellow). Each cell in the matrix represents the correlation of test trials of each image class with training trials of the same class (diagonal) and of other classes (off-diagonal). *Second-order RSA*: First-order RSA test trials are divided into second-order training and test trials. In the next step, we correlate the first-order RSA correlation vector of each second-order test trial with the first-order RSA correlation vectors of all second-order training trials. *Decoding based on RSA-derived values*: The class whose second-order training trials show the highest similarity to that of a given test trial is predicted as that trial’s class. Decoding accuracy (AOC) for each class was quantified by the difference between the fraction of test trials correctly labeled (true positives) to those incorrectly labeled (false positives). **b,** Decoding accuracy for each class within the action sequence learning task (all trials time locked to head turn onsets). The left panels show, for each class, the decoding accuracy as a function of both timepoint along the training trials (y-axes) and that along the test trials (x-axis). The right panels show decoding accuracy averaged along the diagonal (± 2 s on the training time axis), with the shaded regions around each curve depicting the SEM. The horizontal bars (1D) and black contours (2D) in each panel indicate statistically significant decodability (cluster-based permutation test, cluster-forming threshold α = 0.05, overall α = 0.05, 10^6^ permutations, two-sided dependent t-test against zero). In all panels, the average head velocity is depicted in black, with the shaded region indicating the SEM. All time axes are relative to head-turn onset. Panels in **c** show the same as in **b**, but now averaged over all four classes. Both **b** and **c** show that neural activity patterns during action sequence learning can be used to decode the correct image-class on the screen and that such decodability is temporally aligned to the participants’ head turns. Photos: unsplash.com.

Action sequence learning trials were segmented into the four head turn segments of a sequence. Each segment was time-locked to a head turn onset, capturing the participant’s gaze shift toward an image presented on one of the four peripheral screens (exemplary sequence in Fig.1a lower panel, see Methods). Action sequence retrieval trials were segmented in the same way, with each head turn reflecting a self-paced selection of the next head orientation in the sequence (−60°, −30°, 30°, 60° (randomized), exemplary sequence Fig.1b, lower panel).

### Memory for Actions and Contents are Behaviorally Coupled and shows Primacy Effects

Participants (N=36, 24 females, age 24.4 ± 3.7 years) were successfully able to perform both recall tasks (mean and standard derivation of performance for correct reenactment of the full head turn sequence: 0.77 ± 0.16, and correct sequential sorting of images: 0.88 ± 0.18). To further evaluate memory performance, we sorted head turn segments from the action sequence retrieval task as either remembered (i.e., correctly executed) or forgotten (i.e., incorrectly executed). The accuracy for head turns executed to the correct screen position was high for all four positions in the sequence (Fig.1c). However, there was a significant main effect of the sequential position in the head turn sequence on the memory performance (F (3,105) = 11.98, p < 0.001). Specifically, the first head turn of a sequence was remembered most accurately, followed by the second, which was in turn significantly better recalled than the third (Fig.1c, head turn 1 vs. 2: t (35) = 3.26, p = 0.015; head turn 1 vs. 3: t (35) = 5.11, p < 0.001; head turn 1 vs. 4: t (35) = 3.68, head turn 2 vs. 3: t (35) = 3.10, p = 0.023, head turn 2 vs. 4: t (35) = 1.63, p = 0.112, head turn 3 vs. 4: t (35) = −0.94, p = 0.068; all p-values are Bonferroni corrected for multiple comparisons). This main effect of sequential position in the head turn sequence reflects the well-known primacy effect, describing the memory benefit for the first item in list-learning and free recall paradigms (Murdock, 1962). Here, we show that this effect extends to action retrieval, i.e. within a sequence of actions, memory performance is the best corresponding to the first action performed.

In order to assess if the participant’s memory of the action sequences (Fig.1c) were coupled to that of the sequence of images on the screen, we computed the participants’ image sorting performance for correctly executed (i.e. remembered) and incorrectly executed (i.e. forgotten) segments of head turns in the action sequence retrieval task (Fig.1b). We hypothesized that incorrectly executed head-turn segments in the action retrieval task would be associated with incorrect identification of image items in the image sorting task. Indeed, we found that images linked to head-turn segments that were not correctly reenacted during action sequence retrieval were significantly less likely to be placed on their screen in the correct sequential position compared to images associated with successfully retrieved head turns (t (31) = −3.24, p = 0.003, Fig.1d). We thus find behavioral coupling between memory for actions and the memory for contents.

### Action-Locked Decodability of Image Class from Neural Activity

A key goal of our study was to determine if the recall of the head turns in the action sequence retrieval task (Fig.1b) was accompanied by reactivation of neural activity patterns of the associated image class, thus establishing a clear link between action reenactment and the reactivation of visual-stimulus-related neural activity patterns. The neural activity patterns were all time-locked to self-generated motion, i.e., the onset of each head turn. We used a second-order Representational Similarity Analysis (RSA, Fig.2a, see Methods), to compare neural activity patterns between a training set and a test set of trials. The resulting correlation values of the RSA were used to predict the image class of a given head turn segment. Decoding accuracy (hereafter, Accuracy Over Chance, AOC) was calculated as the difference between the ratio of true positives (correct classification of target-image class) and false positives (incorrect classification of non-target image class segments as belonging to the target class; see Methods).

We first assessed our neural-activity decoding technique by training and testing within the sequence learning task, i.e. when the images were displayed on the screen. The decoding accuracy matrices, which plot training segment time points (y-axis) against test segment time points (x-axis) for each class separately, and for the class average, are shown in Fig.2b left panel and Fig.2c upper panel, respectively. A cluster-based permutation test (cluster-forming threshold α = 0.05, overall α = 0.05, 10^6^ permutations, two-sided dependent t-test against zero) on these decoding accuracy matrices revealed a prominent diagonal pattern with significantly elevated decoding accuracy for faces (summed T-values = 271,98, p < 0.001), objects (summed T-values = 179.17, p = 0.013) and houses (summed T-values = 437.17, p < 0.01) as well as for the class average (summed T-values = 751.48, p < 0.001), but not for animals (summed T-values = 30.67, p = 0.61). This diagonal of enhanced decodability shows that decoding accuracy was specifically enhanced when training and test data were temporally aligned through self-initiated action (i.e. gaze shifts). Next, we averaged values along the diagonal of the accuracy matrix using a ±2 time-point window around each training time point. A cluster-based permutation test (cluster-forming threshold α = 0.05, overall α = 0.05, 10^6^ permutations, two-sided dependent t-test against zero) revealed that class identity was significantly decodable in the sequence learning task, with the enhancement in decodability aligning with the head motion (summed T-values = 47.95, p < 0.001, across all classes, Fig.2c lower panel). Again, the correct class was significantly decodable individually for faces (summed T-values = 22.45, p < 0.01), objects (summed T-values = 20.55, p = 0.014) and houses (summed T-values = 41.02, p < 0.001), but not animals (Fig.2b right panel).

### Image Class Decoding Reveals Action-Locked Memory Reactivation During Sequence Retrieval

Having established that our RSA-based decoder could successfully predict the image-class during action sequence learning (where the images were presented on the screens), we used the same technique to address our key question: is action reenactment coupled to neural reactivation of image classes, in the absence of the visual presentation of the images? We used a cross-decoding approach (see Methods for details) in which we trained our RSA-based decoder on segments from the sequence learning task (Fig.1a) and tested it on segments from the action sequence retrieval task, when the participants re-enacted the head turn sequence, with self-paced head turns while only fixation crosses were displayed on all four screens (Fig.1b). Fig.3a left panels and Fig.3b show the decoding accuracies as a function of timepoints along both the training and the test time axes (i.e., similar to Fig.2c upper panel, but now with the test data timepoint being that of the action sequence retrieval task). We find a clear increase in decoding accuracy along the diagonal of the two-dimensional matrix for the majority of image classes (cluster-based permutation test with cluster-forming threshold α = 0.05, overall α = 0.05, 10^6^ permutations, two-sided dependent t-test, tested against zero; decoding accuracy for face: summed T-values = 620.23, p < 0.001; object: summed T-values = 290.01, p < 0.01; house: summed T-values = 495.78, p < 0.001; Fig.3a left panels) and the class average (first cluster: summed T-values = 943.24, p < 0.001, second cluster: summed T-values = 610.47, p < 0.001; Fig.3b). We thus find that the correct images classes are being reactivated from memory as participants re-enact their action sequences, despite the absence of images on the screen. The clear decodability of the correct image class in the associated head-turn segment shows that the reactivation of image-class memory unfolds in a similar temporal sequence that was learnt during action sequence learning.

### Memory Reactivation Tracked by Decoding Accuracy Aligns to Head Motion Velocity

We assessed the symmetrical structure of decoding accuracy across training and test time axes relative to head motion onset, by averaging values along the diagonal of the accuracy matrix using a ±2 time-point window around each training time point. This yielded a one-dimensional time course of decoding accuracy across the time axis of the test dataset for each class (Fig.3a right panels, for each class separately, and Fig.3c, for the average of all classes). A cluster-based permutation test (cluster-forming threshold α = 0.05, overall α = 0.05, 10^6^ permutations, two-sided dependent t-test against zero) revealed that overall class identity was significantly decodable in the action sequence retrieval task (first cluster: summed T-values = 114.02; p < 0.01, second cluster: summed T-values = 37.31, p < 0.01; Fig.3c). Indeed, this cannot be attributed to just one class, as each class was decodable individually (face: summed T-values = 104.02, p < 0.001; object: summed T-values = 34.89, p < 0.01; house: summed T-values = 59.91, p < 0.001 and animal: summed T-values = 37.33, p < 0.001; Fig.3a right panels). We carried out multiple control analyses to find that the significant decodability above cannot be due to an indirect decoding of the head orientations rather than image classes (see Supplementary Material for an extended discussion and Supplementary Figures S.2-4).

Strikingly, the decoding time course closely mirrored the head velocity profile, indicating that class-specific memory reactivation is temporally aligned with self-initiated gaze shifts. Decoding accuracy increased strongly with the current, and less pronounced with thesubsequent head turn (Fig.3c). The increase in accuracy observed with the upcoming head turn can be attributed to the fact that some training and test segments originate from the same sequence and therefore share the same subsequent class (see Supplementary Material). Overall, our cross-decoding results show that neural activity patterns corresponding to image-classes are reactivated sequentially, with clear temporal alignment to the reenactment of the associated head turn.

### Higher Image Class Decodability for Correctly Performed Head Turns

Our behavioral results show a coupling between the memory of the action sequences and the memory of the image sequences, with head turns that were made to the incorrect screens being significantly associated with poorer image memories (Fig.1d). We thus hypothesized that such incorrectly executed head turns would also be associated with lower neural reactivation, and hence lower decodability, when compared to head turns that were correctly executed. We hence investigated whether the decodability of class-specific information differed between correctly (remembered) and incorrectly (forgotten) executed head turns in the action sequence retrieval task (Fig.4). We used the same cross-decoding approach as in Fig.3, to compute decoding accuracies for the remembered and forgotten segments, separately. This analysis was carried out with N=28 participants for whom we had sufficient number of trials in both conditions (see Methods).

**Fig. 3.**
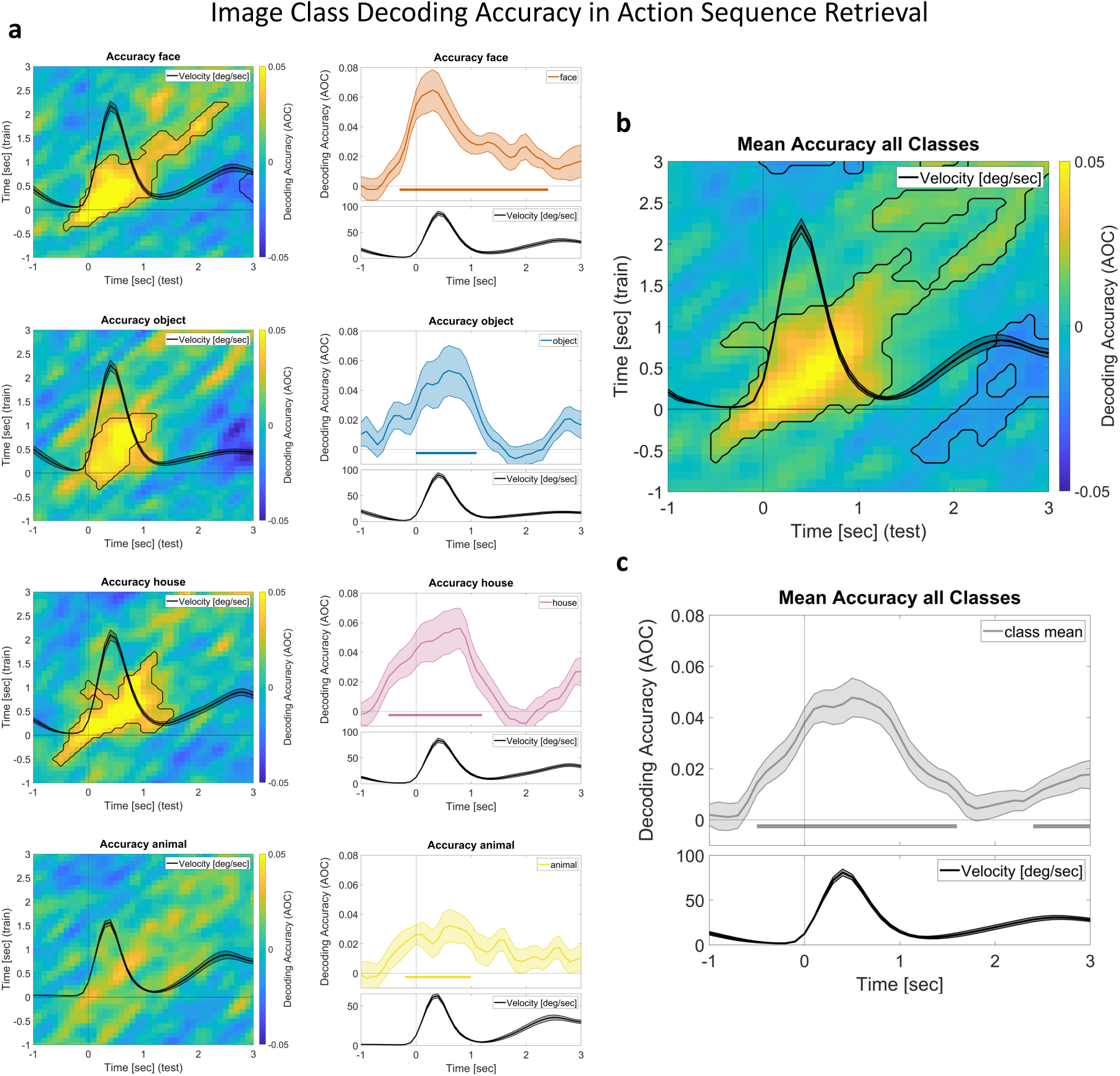
Image-class decoding accuracy in the action sequence retrieval task. The panels in the figure show the outcome of our cross-decoding approach, with head turn segments from the action sequence learning task used as the training set and those from the action sequence retrieval task as the test set. **a**, the left panels show, for each class, the decoding accuracy as a function of both time points in the training (y-axes) and test segments (x-axes). The same is shown in **b**, but for the average across all four classes. **a**, right panels show, for each class, the decoding accuracy averaged along the diagonal (± 2 seconds along the training time axis), with the shaded region indicating the SEM. **c**, shows the same, but now averaged across all four the classes. Significant clusters with decoding accuracy values above chance, identified using a cluster-based permutation test (cluster-forming threshold α = 0.05, overall α = 0.05, 10^6^ permutations, two-sided dependent t-test against zero) are indicated by horizontal bars (1D) and black contours (2D). All trials were time locked to head turn onsets. In all panels, the average head velocity profiles are shown in black, with the shaded region indicating the SEM. The figure shows that image-class memory is reactivated, time-locked to the associated head turns in the learned sequence.

**Fig. 4.**
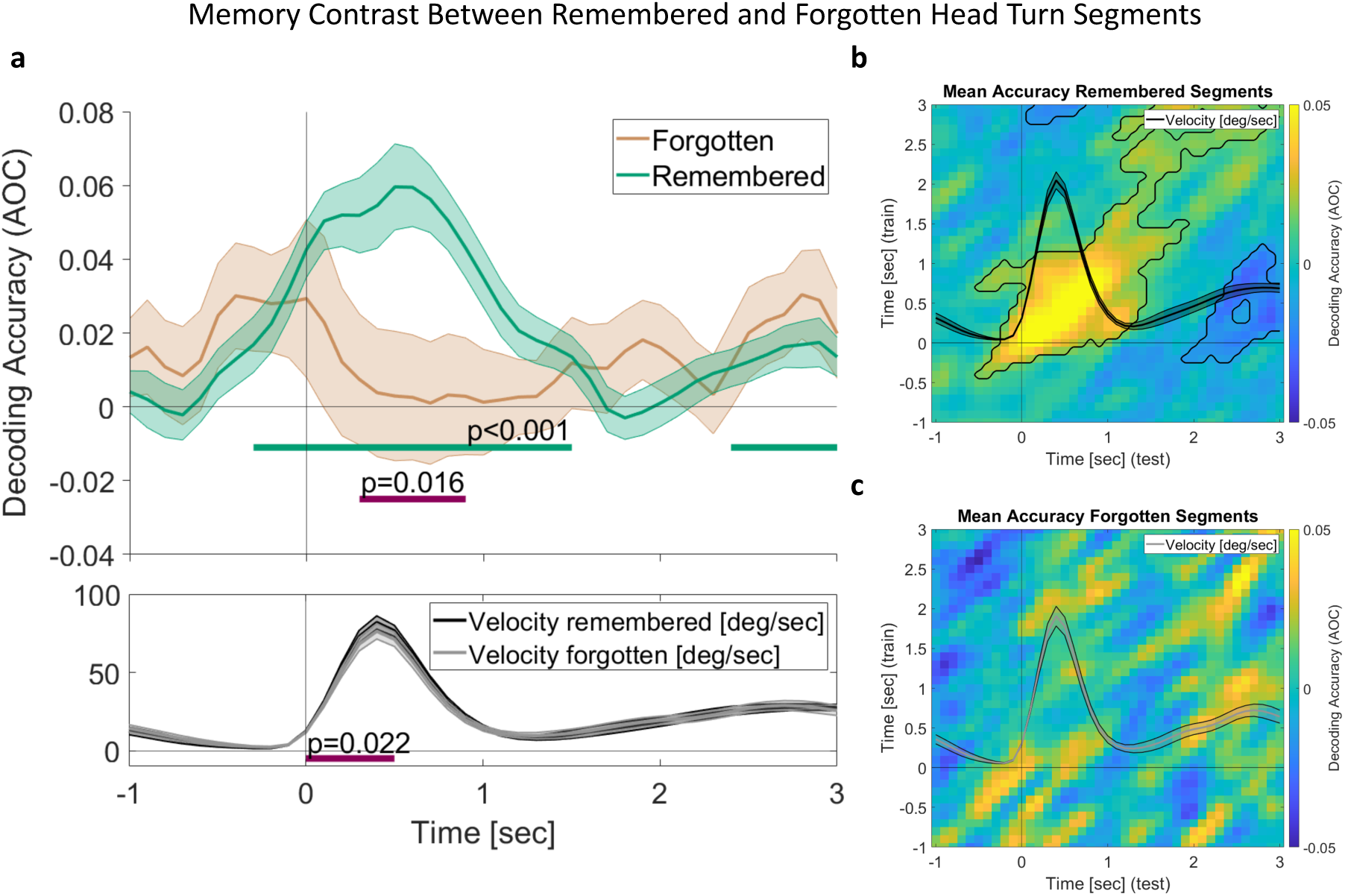
Decoding accuracy for remembered and forgotten head turn segments in action sequence retrieval. **a,** Mean decoding accuracy for remembered (green) and forgotten (brown) head turn segments in the action sequence retrieval task, time-locked to head motion onset. The shaded region around each curve indicates the SEM. Green bars indicate clusters where decoding accuracy for remembered trials was significantly above chance. Magenta bars indicate clusters showing a significant difference in decoding accuracy (upper panel) and velocity (lower panel) between remembered and forgotten trials. **b,c** Decoding accuracy as a function time in training trials (action sequence learning, y-axis) and test trials (action sequence retrieval, x-axis) for the **b** remembered and **c** forgotten head turn segments. Significant clusters were identified using cluster-based permutation tests (cluster-forming threshold α = 0.05, overall α = 0.05, 10^6^ permutations, two-sided dependent t-test against zero) and are indicated by horizontal bars (1D) and black contours (2D). The figure shows that decoding accuracies of associated image classes for remembered head turns are significantly higher than those for forgotten (i.e. incorrectly executed) head turns.

Fig.4a (upper panel) shows the decoding accuracies for the remembered (green curve) and forgotten (brown curve) head turn segments. Decoding accuracy for remembered segments closely followed the head-turn velocity profile (Fig.4a upper panel), with significant cluster that indicates decoding accuracy values significantly above zero (green curve, first cluster: summed T-values = 84.89, p < 0.001; second cluster: summed T-values = 17.8, p = 0.02). The two-dimensional decoding accuracy matrix for the remembered trials (Fig.4b) showed significantly elevated decoding accuracies along the diagonal (positive cluster: summed T-values = 1443,53, p < 0.001), mirroring the temporal dynamics observed in the overall accuracy results of Fig.3. Conversely, decoding accuracy for forgotten segments did not significantly differ from zero at any time point during the action retrieval task (Fig.4a brown graph and Fig.4c). Indeed, a group-level statistical contrast showed that the average decoding accuracy was higher for remembered than for forgotten head-turn segments (cluster-based permutation test with cluster-forming threshold α = 0.05, overall α = 0.05, two-sided dependent-samples t-test, 10^6^ permutations; summed T-values = 16.23, p = 0.016). Note that forgotten segments are identified based on actual head turns (performed to an incorrect screen), just like remembered segments (head turns preformed to the correct screen).

Interestingly, head velocity in remembered trials was also significantly higher than in forgotten trials (cluster-based permutation test with cluster-forming threshold α = 0.05, overall α = 0.05, two-sided dependent-samples t-test, 10^6^ permutations; summed T-values = 14.16, p = 0.022; Fig.4a, lower panel, purple significance bar), reflecting differences in behavior associated with memory. Head-turns for remembered segments are thus executed faster and are associated with the reactivation of the correct image memory, in contrast to forgotten segments, as revealed by elevated decoding accuracies.

## Discussion

Employing a paradigm where self-initiated gaze shifts, i.e., the coordinated movement of head and eyes, functions as behavioral readout of memory sequences, we show that these actions can support the temporal alignment of neural activity that indicates the reactivation of episodic memory content.

### Gaze Shifts Structure Reactivation of Sequential Episodic Memory Content

During memory retrieval, participants actively recalled action sequences through a specific self-paced sequence of gaze shifts. Employing representational similarity analyses, we show that the neural patterns of images previously associated with the action sequence were reactivated with corresponding gaze shifts. Image class could be decoded when training and testing on EEG activity during sequence learning, and, crucially, also when training on the learning phase and testing on the retrieval phase, in which no images were presented on the screens. Decoding of the image class was possible when both training and testing data were time-locked to head turn onsets, suggesting that self-initiated actions are integrated with the encoding and reactivation of an episodic memory trace, and can serve as a reliable behavioral marker of the temporal unfolding of episodic memories.

The accuracy of image class decoding aligned with head velocity, such that it increased with movement onset and declined as the head stabilized. This indicates that locking to head turn onset temporally aligns brain activity, leading to highest decoding accuracy at corresponding time points of training and test trials. The temporal evolution of image-class-related brain activity following head-turn onset during sequence learning parallels findings from human object recognition, where object category decoding was found to be possible within a time window of ∼50–900 ms after stimulus onset (Carlson et al., 2011, 2013; Cichy et al., 2014). The source of these decodable signals has been suggested to be highly transient neural activity that propagates along the ventral visual stream to process different image properties at successive time points (Cichy et al., 2014, 2016). At the same time, periods of temporal generalization across neighboring time points during the first second of image perception produce a broadened diagonal of increased similarity (Cichy et al., 2014; King & Dehaene, 2014), consistent with the temporal evolution pattern we find in our own data. During memory recall, we too observed a window of increased image class decoding accuracy along a broader diagonal, peaking ∼500 ms after head turn onset. Since the decision of where to move the head next has to precede the increase in head turn velocity, we assume that action production related processes help structuring the neural reinstatement of episodic memory content. Together, our findings align with prior evidence showing neural representations at early stages of encoding, and reactivation patterns occurring ∼500 ms after cue onset during retrieval (Jafarpour et al., 2014; Lu et al., 2015; Staresina & Wimber, 2019). In our study, memory reactivation during retrieval mirrors the temporal dynamics of brain activity during encoding, with self-initiated action serving as a shared temporal anchor across both phases.

Head turns during action sequence retrieval served as behavioral markers of memory recall: image class decoding performance was significantly higher for correct head turns than for incorrect ones. This difference and the lack of a significant reactivation of the correct image in incorrectly executed head turns, indicate forgotten or confused images. This shows that the accuracy of actions predicts memory reactivation, reflecting the interdependence between action and image memory: when one is forgotten, the other tends to be as well. Successful retrieval was thus associated with stronger, more class-specific neural reinstatement, consistent with prior work linking accurate memory retrieval to reactivation of memory-relevant neural representations (for reviews, see Rugg et al., 2008; Schreiner & Staudigl, 2020). Together, this pattern of results indicates that the accuracy of performing an action reflects the fidelity of the underlying memory trace.

### Behavioral Evidence for Action Sequences

Beyond the neural evidence for memory reactivation, our paradigm enabled a direct behavioral read-out of memory sequences which also indicated that the episodic memories in our task are structured around action sequences. Participants successfully recalled gaze shift sequences from the learning phase and subsequently reconstructed the associated sequence of images. Importantly, the behavioral outcomes reflected a coupling of actions and contents in memory: sequential position and location of images associated with incorrectly executed gaze shifts were more likely to be forgotten as compared to those associated with correctly executed shifts. This correspondence between action sequences and memory contents indicates that both are integrated into a shared episodic memory trace. The behavioral findings thus complement the representational similarity analyses in showing that successful recall of an action predicts the successful retrieval of the associated content.

The link between action and underlying memory traces becomes especially evident when relating action velocity to memory performance: the velocity of correctly performed head turns during the recall of action sequences was systematically higher compared to that of incorrectly performed head turns. While the mechanistic relationship between the latency of reactions and memory performance in memory tasks is complex, the two are nonetheless typically found to be correlated (Kahana & Loftus, 1999). Interestingly, in a study design with self-paced actions, we find that it is the speed of the action itself, that is predictive of memory performance. At the same time, reaction times and memory confidence ratings have been found to be highly correlated, which is thought to indicate a shared relationship to memory fidelity (Weidemann & Kahana, 2016). It is thus possible that the speed of gaze shifts in our action sequence recall acts as an indicator of successful memory retrieval and retrieval confidence.

Memory performance in our task also depended on the serial position within the learned sequence, with the first gaze position being remembered more accurately than later positions. This resembles the well-established primacy memory effect, in which items at the start of a list are recalled better than items presented later (Kahana, 2020; Murdock, 1962). In classical accounts, primacy effects have been attributed to rehearsal effects, greater attentional allocation, or stronger contextual binding of early list items (Atkinson & Shiffrin, 1968; Brodie & Murdock, 1977; Howard & Kahana, 2002; Tan & Ward, 2000). In our paradigm, the first gaze shift may play an important role as initial temporal anchor of the sequential memory trace, from which the remaining elements of a sequence unfold. This interpretation can be related to the temporal context model (Howard & Kahana, 2002), which proposes that experiences occur within a gradually changing temporal context. Memories are associated with the current contextual state, and using the particular context as a cue helps retrieving the associated memories (Polyn et al., 2009). To avoid constant cue overload, each new experience is associated with a distinct temporal context. Relating this account to our findings, self-generated actions - and in particular the first action within a sequence - could contribute to initiating a new contextual state that distinguishes one sequence from the next. From this perspective, actions could serve as internally generated event boundaries (see below) that organize experiences into distinct memory episodes (DuBrow & Davachi, 2013; Zacks & Swallow, 2007).

The sequential nature of our findings connects them to the literature on replay (see, e.g. Genzel et al., 2020; Schreiner & Staudigl, 2020; Schuck et al., 2026). Replay can be observed when experience-induced sequences of neuronal activity reoccur. The classic findings in rodents demonstrated replay by showing sequential reactivation of place-cell firing patterns during periods of rest and sleep (Davidson et al., 2009; Diba & Buzsáki, 2007; Foster & Wilson, 2006). In humans, replay has been described non-invasively (Y. Liu et al., 2019; Michelmann et al., 2019; Schuck & Niv, 2019). Whether the sequential reactivation observed here – expressed through recalled action sequences accompanied by neural reactivation at each action – should be considered a form of replay remains an open question. One speculative interpretation of the role of actions in replay is that self-generated actions provide temporal anchors allowing the remaining elements of a sequence to unfold from these anchor points.

### Limitations

Because our paradigm allowed participants to freely move their head and eyes, two movements that are strongly reflected in brain activity and as artefacts in scalp EEG, we implemented several design choices and control analysis to rule out confounds from head-movement-related signals. Critically, our image class decoding was based on training and testing on image class, not head turns: image classes were randomized across all head orientations, preventing any explicit image–orientation association. In addition, two control analyses further excluded residual confounds: decoding performance was unrelated to any residual image-class–head-orientation bias within participants (Supplementary Material) and including head-turn localizer trials (head turns with fixation crosses only, no images) in the training data left decoding performance unchanged (Supplementary Material), confirming that decoding was not driven by head-turn-specific patterns. Together, these analyses support genuine image memory decoding in our main analyses, and speak against head-movement-related activity or artefacts as confounding factors.

### Action and Memory: A Coupled System

Our finding that gaze shifts align neural representations of perception with the reactivation of episodic memory content suggests that gaze shifts act as temporal structuring elements, potentially providing a scaffold for internal neural rhythms that organize episodic representations (Buzsáki, 2006; Buzsáki & Tingley, 2018).

This builds on a growing body of work on active sensing (Ahissar & Arieli, 2001; Kleinfeld et al., 2006; Schroeder et al., 2010), and memory, which has shown that eye movements are closely linked to both encoding and retrieval of visual information (Barker et al., 2026; Fehlmann et al., 2020; Johansson & Johansson, 2014; Kinjo et al., 2020; Lucas et al., 2023; Paeng & Kim, 2024; Wu et al., 2025). Gaze reinstatement, the recurrence of viewing patterns during recall that mirror those during encoding, is closely tied to memory retrieval (Henin et al., 2025; Johansson et al., 2022; Kragel & Voss, 2021; Wynn et al., 2019) and to neural reinstatement (Johansson & Johansson, 2014; Nau et al., 2025; Wynn et al., 2022). Our study extends this framework by showing that gaze shifts, incorporating head movements as the other core component of natural gaze, may structure the sequential reactivation of an entire memory episode.

The link between action and memory reactivation may be mediated by interactions between gaze-shift components (saccades and head movements) and brain dynamics in memory-critical regions such as the medial temporal lobe (MTL) (Kragel & Voss, 2022). In non-human primates, saccades induce low-frequency phase resets in the MTL (Jutras et al., 2013). In human MTL structures, they similarly align low-frequency phase and elicit ERPs (Hoffman et al., 2013; Katz et al., 2020, 2022; Staudigl et al., 2022), coordinating with and structuring memory-related processing (Leszczynski & Schroeder, 2019; Staudigl et al., 2017). Head movements also interact with MTL dynamics: head-gaze shifts reset hippocampal theta in marmosets (Piza et al., 2024) and MTL theta power increases during human head scanning behavior (Aghajan et al., 2017). Parahippocampal activity tracks heading direction (Griffiths et al., 2024) and MTL activity during head turns further segments navigational trajectories in real-world and imagined navigation (Seeber et al., 2025). While our study is agnostic to how MTL regions might contribute to the unfolding of sequential memory reactivation, future studies could investigate how gaze-related MTL activity contributes to memory processes.

Self-initiated actions might support parsing continuous experience into discrete episodic units, similar to event segmentation in ongoing perceptual processing (Zacks & Swallow, 2007), a process crucial for memory and planning (Kurby & Zacks, 2008). Event boundaries are typically studied in visual streams like movie clips, where changes in visual input, scene dynamics, and spatiotemporal context signal transitions between events (Zacks et al., 2009), linked to cortical (Lee & Chen, 2024) and hippocampal activity (Silva et al., 2025). Our study raises the question if actions themselves might also serve as event boundaries. Recent work showed that hippocampal population activity in monkeys shifts abruptly at behavioral events, parsing activity into discrete ensemble codes (Rueckemann et al., 2025). More directly related to our findings, eye movements have been found to reflect event segmentation, linking boundary perception to oculomotor parameters such as saccade speed and pupil size (Li et al., 2025; Smith et al., 2024). Head and eye movements operate on different timescales, with multiple saccades occurring within single head movements (Fang et al., 2015). Whether these actions scaffold memory reactivation through hierarchical event boundaries operating simultaneously across multiple temporal scales (Kurby & Zacks, 2008), needs to be investigated directly in future studies.

Additional to the suggested temporal scaffolding, actions also contribute spatial cues to memory traces. In virtual reality tasks, spatial-context reactivation precedes episodic recall (Herweg et al., 2020). The quality of spatial context representations further predicts cortical reinstatement of associated objects (Masís-Obando et al., 2026). Action-derived spatial information, such as gaze-shift self-motion cues, may thus act as a contextual feature (Polyn & Kahana, 2008). By introducing distinct spatial contexts for each image in our sequences, gaze shifts could support context-dependent encoding by disambiguating similar contents within and across sequences through unique spatial contexts.

This study focused on the most frequently performed exploratory behavior in humans: head and eye movement driven gaze shifts. However, other forms of action may also contribute to memory processes as proposed in the enactment effect (Cohen, 1981; Engelkamp & Krumnacker, 1980). This effect demonstrates enhanced memory for action-involving compared to passively perceived contents (Cook et al., 2010; Engelkamp & Cohen, 1991; Macedonia, 2014). Various forms of motor involvement may thus enrich contextual encoding, potentially by integrating a motor trace into memories (Albouy et al., 2015; Dolfen et al., 2024; Macedonia et al., 2011).

## Conclusion

Our findings reveal a tight coupling between self-initiated actions and episodic memory reactivation: first, gaze shifts align brain activity of perception and recall, enabling decoding of memory reactivation; second, correctly recalled gaze shifts are accompanied by the reinstatement of the correct image memory, while misperformed gaze shifts are not. This suggests that action and memory content are not only aligned in time but entwined with one another. This coupling highlights the potential for self-generated movements to act as internal anchors that structure encoding and retrieval, organizing memory into action-aligned segments that may facilitate the reconstruction of complex experiences. This supports an active sensing account of memory, in which movement is not a passive byproduct but actively interacts with the brain dynamics underlying memory processes. Future studies could attempt to directly manipulate actions during memory processes and ask whether disrupting an action impairs the reactivation of associated memory contents, and whether disrupting memory reactivation in turn destabilizes the execution of the associated action. Taken together, incorporating self-initiated action into memory research offers a more ecologically grounded account of episodic memory, with implications for understanding memory during natural behaviors outside the laboratory. Our findings suggest actions are not merely a byproduct of remembering but structure the encoding and reactivation of memory traces.

## Methods

### Participants

We collected data from 39 healthy participants, ensuring that none had any neurological or psychiatric conditions. Due to technical issues with the EEG and motion tracking systems, data from 3 participants had to be excluded, resulting in a final sample of 36 participants comprising 12 males and 24 females, with a mean of age 24.4 ± 3.7 years (mean ± SD). Participants were recruited via student information platforms at LMU Munich and were compensated either with course credit or a monetary reward. All participants provided written informed consent after receiving detailed information about the study. The study was approved by the local ethics committee.

### Paradigm

This study was part of a larger experiment. We describe below only parts of the experiment relevant to this study. The experimental setup featured a five-screen configuration surrounding the participant who was seated in a chair facing the central screen. Four screens were flanking the central screen at −30°, −60°, 30° and 60° angles (Fig.1a,b upper panel). Participants performed memory tasks consisting of three identical blocks, each block comprising the learning and immediate retrieval of eight unique sequences. At the end of each block, memory performance for these eight sequences was tested in a retrieval test. This procedure resulted in a total of 24 learned and recalled sequences per participant.

Each sequence consisted of four images, each of which was consecutively presented for 3 seconds (±0.1 s jitter) on one of the peripheral screens. The four images were such that there was one image from each of the four classes (faces, objects, houses, animals). Further, each sequence of four images had an associated unique sequence of head turns. The paring of image class and sequential position was randomly generated and balanced across all 24 sequences, to make sure each class was presented the same number of times at each position in the sequence. Each sequence was preceded by an auditory cue, a distinct German verb that was played aloud on speakers. Participants were instructed to shift their gaze, by turning their head toward a screen whenever it displayed an image and to remain in this position for the duration of presentation. They were asked to memorize both the sequence of images, and the associated screen positions, ultimately constructing a spatiotemporal sequence that integrated information from both the visual stimuli and the corresponding head movements (action sequence learning). To facilitate memorization of the sequence, participants were instructed to construct a narrative around the sequence that began by using the verb as a cue. After seeing the image sequence, they verbalized this narrative, while fixation crosses were presented on all screens. Finally, the images of the sequence were repeated on the screens and participants again turned their head accordingly, confirming the correctness of their memorized story. The initial presentation of the image sequence and its repetition constitute the action sequence learning trials. Following the learning of one sequence, participants were asked to immediately perform three retrieval tasks, identical to the tasks in the subsequent episodic retrieval test described below.

After the learning and immediate retrieval of the eight sequences, participants completed a three-minute-long distractor task. Subsequently, they proceeded to the sequence retrieval test. To cue the retrieval of a sequence within each block of eight, participants were presented with one of the verb-sounds that were uniquely associated with a sequence during the action sequence learning phase. Note that the order of presentation of the eight sequences was randomized and was thus different in retrieval and learning. Participants performed three retrieval tasks: 1) retrieving the cued sequence while looking at the center screen, 2) re-performing the sequence of head turns while recalling the full episode (action sequence retrieval task, Fig.1b), and 3) selecting and placing the images in the correct order on grey spatial placeholders representing the screen setup, displayed on the center screen (image sequence sorting task, Fig.1d). After retrieving all eight sequences of a block, the action sequence learning phase of the next block started.

Preceding the three memory task blocks, participants performed a set of non-memory localizer tasks, including a head turn localizer task used for control analysis. Participants were instructed to follow a fixation cross as it appeared sequentially on the four peripheral screens. Upon each appearance, participants moved their head to align their gaze with the cross and maintained fixation until the next location was cued. Finally, they returned their gaze to the central fixation cross. Essentially, participants performed all sequences of head turns that were part of the later memory tasks.

### Data Collection and Processing

All data processing and statistical analyses were performed in MATLAB (R2020a) using the FieldTrip environment (Oostenveld et al., 2011).

#### EEG Acquisition and Preprocessing

We used an EEG system (EEGo, ANT Neuro, Enschede, Netherlands) with a 10/10 system layout, consisting of 66 Ag/AgCl electrodes, including both mastoids and the inion. Additionally, we recorded two electrooculography (EOG) channels. CPz served as the reference and AFz as the ground. Impedance values of all electrodes were required to be <20 ohms for the experiment to begin. The sampling rate was set to 1000 Hz. EEG data acquisition was conducted using ANT Neuro’s LE-200 eego software (version 1.8.2, Build 8.2.26.45751). Electrodes were secured using both an EEG cap and an additional flexible mesh bandage to prevent electrode movement.

We used 63 final channels from our electrode layout, excluding the mastoid electrodes, the inion, and the two EOG channels from all subsequent analyses. The data were band-stop filtered to remove power-line artefacts (49-51Hz, 99-101 Hz; 149-151 Hz; 199-201 Hz; Butterworth IIR) and re-referenced to the common average. Visual inspection of the data was conducted to identify and remove any bad channels. The data were then epoched into 5 s intervals, and Independent Component Analysis (ICA) was applied to these epochs. Components for all epochs were visually inspected, and those corresponding to typical artifacts such as blinks, muscles and heartbeat were discarded. Subsequently, the data were segmented into trials according to the task structure of the experimental paradigm. For each trial, all channels were visually inspected for artifacts, and trials with contaminated channels were excluded. As a final step, the activity from discarded channels was interpolated based on the activity of neighboring channels.

#### Head Motion Tracking

Head rotations were recorded using a Polhemus Liberty system. One motion sensor was affixed to the EEG cap between electrodes AFz and Fz and another one around 18 cm posterior to the first at the back of the head, while the reference sensor device was placed approximately 75 cm behind and to the right of the participant. The system’s proprietary software continuously tracked the position of the two sensors relative to the reference at a sampling rate of 240 Hz. In post-processing, the coordinate data were epoched around the onset of the first visual stimulus in each task, either image onset or fixation cross onset, respectively, aligned with the segmentation used for EEG data. Head angle was extracted from the Euler angle output, using the rotation in the horizontal plane. The resulting head orientation signal was visually inspected for artifacts, and any trial containing physiologically implausible changes in head angle was excluded. Finally, the data were upsampled to match the 1000 Hz sampling rate of the EEG recordings. We computed head rotation velocity by taking the absolute value of the first temporal derivative of the head angle signal, dividing the difference between consecutive angle samples by the sampling interval (head angle and velocity were downsampled to 100 Hz for the plot in Fig.1a,b lower panels). Head motion velocity was subsequently used for trial segmentation and head-motion-locked analyses. Since we used head motion as the temporal marker of self-initiated gaze shifts, “head turn” is used as synonym to gaze shift including head and eye movement towards a screen.

#### Head Turn Segmentation

Trials in both tasks involving head turns, i.e. the action sequence learning task and the action sequence retrieval task, were segmented into four distinct epochs, time-locked to the onset of each head turn to a flanking screen (Fig.1a,b lower panels). This segmentation accounted for both inter-subject and inter-trial variability in head motion dynamics. To identify the onsets of head turns, we first computed the absolute head motion velocity across each trial, downsampled to 100 Hz. We then estimated a baseline mean velocity using all timepoints for which the velocity values were below three standard deviations of the velocity across the entire trial. This baseline mean velocity served as a threshold for detecting significant motion events, corresponding to the four head turns. Timepoints exceeding this threshold were marked as candidate onsets (and offsets) of head turns. These candidate onsets and offsets were used to identify the time periods when the gaze was oriented towards one of the four screens. The four longest of such segments were treated as belonging to the four head turns of interest. We performed additional visual inspection to ensure that any additional brief head movement was not mistakenly identified as a head turn of interest. For all our analyses, we included head turn segments only from trials in which all four head turns were clearly identifiable. This criterion was necessary because if fewer than four head turns could be identified in the motion trace, it would be impossible to determine which head turn corresponded to which position in the sequence. Finally, the four turns were ordered by relative amplitude and direction (largest-right, largest-left, smaller-right, smaller-left) and assigned to the corresponding screen positions (−60°, 60°, −30°, and 30°, respectively). A head turn was considered correct if it was directed toward the expected screen location corresponding to a given position in the learned sequence.

### Data Analyses

#### Behavioral Data Analyses

Repeated-measures models were fit using the Matlab function *fitrm*, followed by repeated-measures ANOVA with the function *ranova*. Post-hoc pairwise comparisons were conducted using paired t-tests with Bonferroni correction to control for multiple comparisons.

#### Representational Similarity Analysis Based Class Decoding

We employed Representational Similarity Analysis (RSA) (Kriegeskorte et al., 2008) to examine the similarity of neural activity within the action sequence learning task, as well as between action sequence learning and action sequence retrieval. We implemented a channel-wise first-order RSA (see *first-order RSA* below), followed by a second-order RSA approach. Specifically, for each trial of a class, we used the temporal correlation patterns between that trial and trials of all other classes (including the same class), derived from the first-order RSA, as a representational signature. These first-order representations were then used as input for the second-order RSA, allowing us to compare the similarity matrices themselves rather than relying directly on neural activation patterns (see *Second-order RSA* below). This approach enabled an assessment of class-specific representational similarity between different tasks. The resulting similarity values thus formed the base for the image class decoding.

Below, we describe the first- and second-order RSA and decoding approach, exemplarily for the within action sequence learning task. Within this task, we used a bootstrapping procedure, applied on the whole analysis pipeline (first- and second-order RSA and subsequent decoding). We used a total of 50 bootstrap iterations. For each bootstrap iteration, we randomly split the total number of trials into independent training and test sets. For each set, we balanced the number of trials for all four classes. For our cross-task decoding analysis (action sequence learning task as training set and action sequence retrieval task as test set), the procedure was identical, except that here, we compare the within similarity of action sequence learning trials to the similarity of learning and retrieval trials at the second-order RSA level (see section *Within and Across Task RSA Based Class Decoding* and Fig.5).

**Fig. 5.**
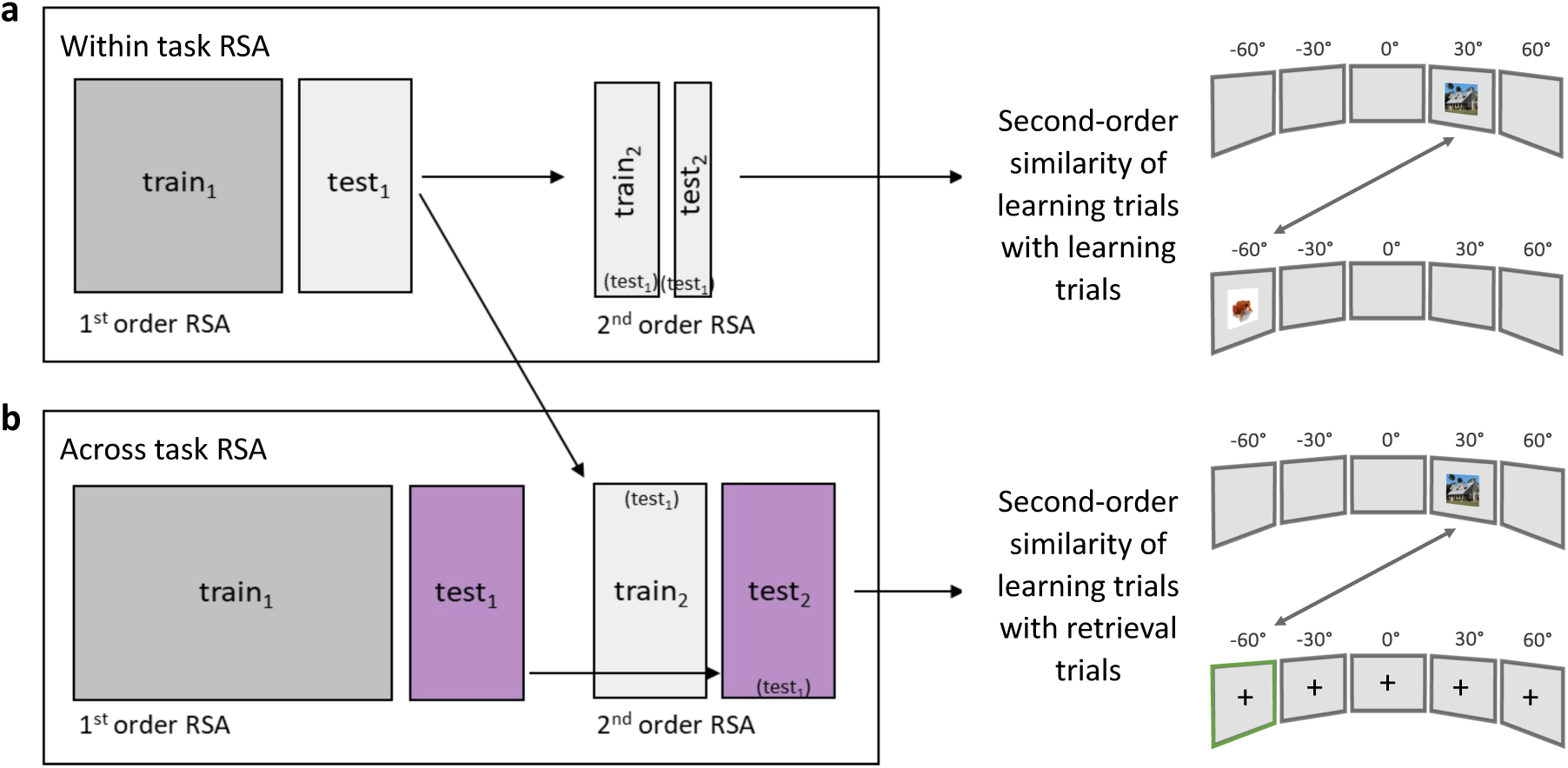
Schematic illustration of the within-task and across-task RSA design. **a,** For the *within-*task *RSA*, we first perform a first-order RSA by training on two-thirds (dark grey, set train1) of the action sequence learning data and testing on the remaining one-third (light grey, set test1). We then further split the test set (two-thirds for training (light grey, set train2) and one-third for testing (light grey, set test2) to compute a second-order RSA on the resulting correlational values. This allowed us to assess the similarity of trials within the learning task. **b,** For the *across*-task RSA, we use the full training set (action sequence learning task, dark grey, set train1) and evaluate it on a different test set (action sequence retrieval task, purple, set test1). We then compute a second-order RSA between the first-order RSA results of the *within*-task (light grey, set train2) and *across*-task (purple, set test2). This allowed us to assess the similarity of learning trials and retrieval trials. Photos: unsplash.com.

##### First-order RSA

We downsampled the data to 100 Hz and segmented it into overlapping temporal bins (hereafter, *ttest1/train1*) of 400 ms duration, with 100 ms steps, within the time range of interest (−1-3 s). We split the data into independent training (hereafter, *train1*) and test (hereafter*, test1*) trials. For each test trial (in set *test1*), we computed the Spearman rank-order correlation with all training trials (in set *train1*). We did that for each channel separately, correlating the 400 ms time bins of training and test trials (i.e. *ttest1* and *ttrain1*). We next computed the mean correlation across all training trials of a given class. This procedure yielded a correlation tensor of the dimensions *channel × training class × test time × training time* for each test trial.

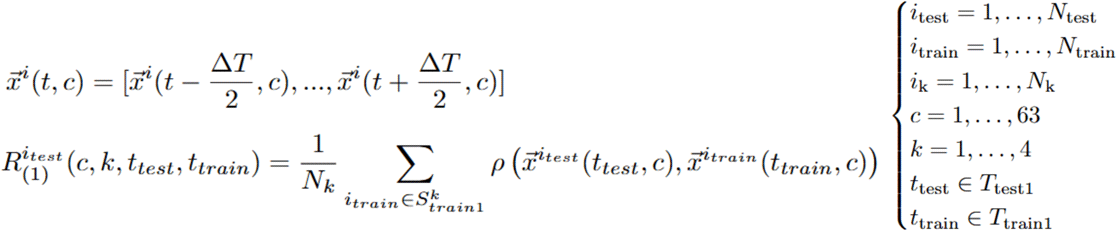

**First-order RSA:** *x^i^(t,c)* is the measured data vector of trial *i* at time bin *t* for EEG channel *c. R ^itest^(1) (c,k,ttest,ttrain)* is the output tensor of Spearman rank-order correlation for test trial *itest* with dimensions channel (indexed by *c)*, training class (indexed by *k)*, test time bin (indexed by *ttest),* and training time bin (index by *ttrain.).* The other symbols in the expression above are as follows. *Ntest1*: Number of test trials; *Ntrain1*: Number of training trials; *Nk*: Number of training trials of class *k*; *S^k^train1*: Set of training trials of class *k*; *Ttest1/train1*: Set of timepoints that mark the center of each time bin (length *ΔT* = 400 ms).

##### Second-order RSA

The output tensors from the first-order RSA (dimensions *channel × training class × test time × training time*) for each trial in the former test set, *test1*, subtracted by the mean correlation across all training classes, served as the input to the second-order RSA. All former test trials (*test1*) were now again split into two subsets: one-third used as the second-order test set (hereafter, *test2*) and two-thirds as the second-order training set (hereafter, *train2*). We again segmented the data along the training and test time into overlapping 400 ms time bins, advancing in 100 ms steps.

For each trial in the new test set*, test2*, we computed time-bin by time-bin Spearman rank-order correlations with all trials in the new training set, *train2*. We correlated the concatenated feature dimensions*: channel × class of the first-order RSA training set (train1) × time-bin size (400 ms) × time of the first-order RSA training set (train1).* We subsequently averaged across all training trials (in set *train2)* of a given class. This resulted in a second-order similarity tensor for each trial in the test dataset, *test2*, with dimensions *class × test time × training time*.

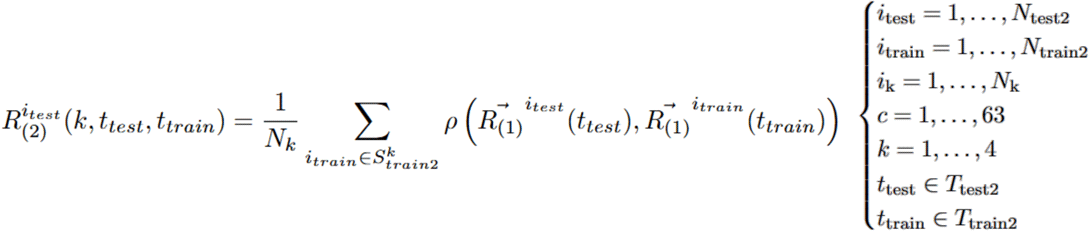

**Second-order RSA:** *R^itest^(2)(k,ttest, ttrain)* is the Spearman rank-order correlation output tensor for test trial *itest*, with the dimensions training class (indexed by *k)*, test time bin (indexed by *ttest),* and training time bin (indexed by *ttrain)*. The other symbols in the expression above are as follows. *Ntest2*: Number of test trials. *Ntrain2*: Number of training trials. *Nk*: Number of training trials of class *k*. *R̅^i^(1) (t)*: Vector of first-order RSA tensor of trial itest and itrain (concatenated dimensions: channel, training class (1^st^ order RSA), time bin (centered at time points *Ttest2/train2*), training time (1^st^ order RSA)), obtained after subtracting the mean across all training classes. *S^k^train2*: Set of training trials of class *k*; *Ttest2/train2*: Set of timepoints that mark the center of each time bin (length *ΔT* = 400ms).

##### Decoding

For the decoding we used the second-order similarity tensor of each test trial in the set *test2*. For every sample in the similarity tensor, indexed by a specific combination of test time point × training time point, we identified the training class that had the highest correlation value. This class was considered the predicted class for that sample in the given test trial. To quantify decoding performance, we computed, for each time sample, the proportion of trials in which the predicted class matched the true class label of the current trial (i.e. class of the image presented on the screen during leaning). To control for false positives, we subtracted the proportion of trials that did not belong to the true class but was still (incorrectly) labeled as the same class label. For example, for the decoding accuracy of houses, we computed the proportion of house trials that were correctly decoded as “house” minus the proportion of non-house trials that were incorrectly classified as “house”. The resulting value thus represents the decoding accuracy for each class, corrected for class specific chance-level classification (true positives rate - false positives rate). To compute decoding accuracy in the action sequence retrieval task, where no visual image was presented on the screen during the recall, we assigned the true class label based on the sequential positions of images during the sequence learning task. For example, for the second head turn of a specific sequence, we used the class label of the image, presented at the second position during sequence learning. Finally, we applied a Gaussian smoothing kernel to all time × time accuracy matrices (*kernel size* = 0.3 s x 0.3 s, *σ* = 0.1 s).

#### Within and Across Task RSA Based Class Decoding

As mentioned above, we performed within task decoding in the action sequence learning task as well as across task decoding, using action sequence learning trials as training and action sequence retrieval trials as test data.

As described in the previous section, for the within-task RSA and decoding, we computed the first-order RSA by training on two-thirds of the data and testing on the remaining one-third. We then performed a second-order RSA, by further splitting the test set (two-thirds for training and one-third for testing, Fig.5a). The resulting correlation values were used for the within task decoding results in Fig.2. To avoid biases due to different trial numbers across classes, we applied a subsampling procedure, where the number of trials per class was equalized by randomly excluding trials (50 bootstrap iterations, random resampling). The final decoding accuracy was then averaged across all 50 bootstraps.

For the across-task RSA, we implemented three steps: First, we performed an *across* task first-order RSA, with all action sequence learning trials as training set and all action sequence retrieval trials as test set. This captured the similarity of learning trials to retrieval trials (for all class and timepoint combinations) and served as the test data for the second-order RSA (Fig.5b). Second, we retrieved the results of the *within* sequence learning first-order RSA (see above and Fig.5b). This captured the similarity of learning trials to learning trials (for all class and timepoint combinations) and served as the training data for the second-order RSA. Finally, we computed the second-order RSA between training and test data. The resulting correlation values were used for the across task decoding results in Fig.3.

#### Accounting for Different Trial Numbers in Memory Contrast

The number of remembered and forgotten head turn segments were, in general, different. In order to ensure comparable variance in decoding accuracy between the two conditions, we used subsampling to equalize, for each participant, the trial count across the two conditions. This was done prior to computing decoding accuracy and repeated multiple times to obtain estimates of the decoding accuracies for remembered and forgotten segments. For each condition and each iteration, decoding accuracy was only computed if at least one head turn-segment per class was present. Bootstrapping continued until 20,000 valid accuracy estimates were obtained per condition. Participants who did not meet these criteria, or for whom sufficient valid bootstrap samples could not be obtained, were excluded from group-level analyses for the memory contrast. This resulted in N=28 participants (out of 36) for the action sequence retrieval memory contrast.

## Acknowledgements

This work was supported by the European Research Council (ERC, ERC-STG Starting Grant 802681 to T. Staudigl). We thank all student assistants who supported the data collection, and all participants for making this study possible. Photos used for illustrative purposes in the figures (Fig.1,2,5) and video (S.1) were obtained from a source of freely-usable images (Unsplash, n.d.); we thank the following photographers: male face: Mitchell Griest; female face: Ionela Mat; house, brown: Johnson; house, grey: Sieuwert Otterloo; elephant: Kaffeebart; frog: Smithsonian; clock: Adriano Pucciarelli; rubik’s cube: Volodymyr Hryshchenko.

## Author Contributions

Conceptualization: TS, JS. Methodology: TS, JS. Investigation: JS. Formal analysis: JS, AC. Writing – original draft: JS, AC, TS. Writing – review & editing: TS, AC, JS. Supervision: TS, AC. Funding acquisition: TS.

## Competing Interests

The authors declare no competing interests.

## Notes

### Competing Interest Statement

The authors have declared no competing interest.

