## Supplementary figures and images for "Action Sequences Structure Human Memory Reactivation"

### Supplementary_Materials_S1

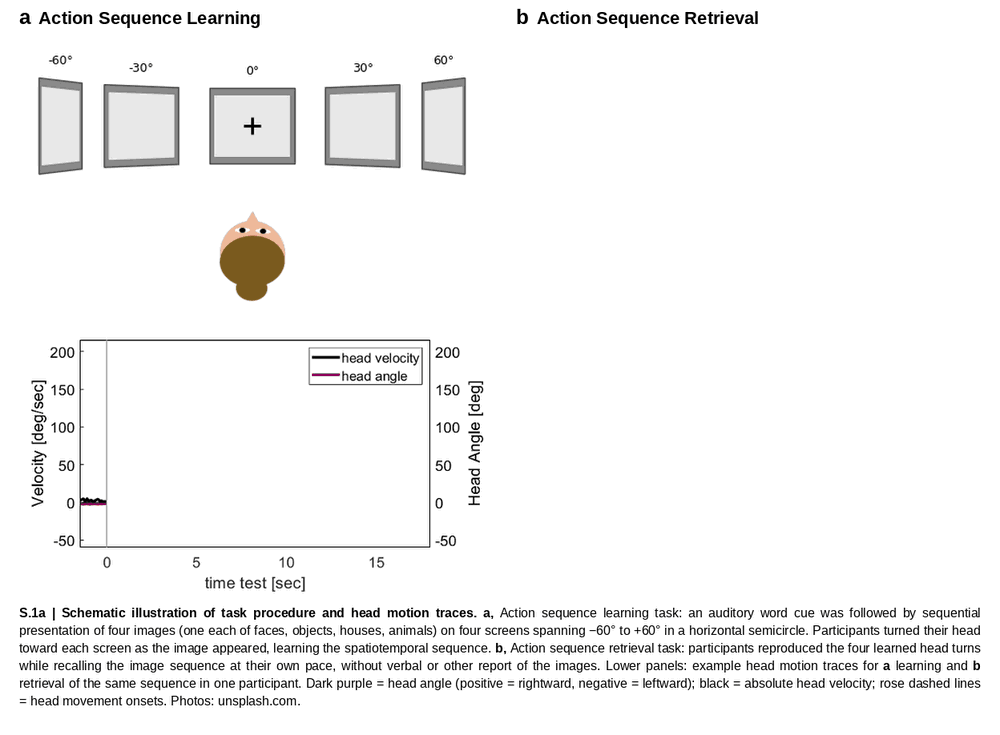
